# Searching the Druggable Genome using Large Language Models

**DOI:** 10.64898/2026.01.18.700012

**Authors:** Lars Schimmelpfennig, Matthew Cannon, Quentin Cody, Joshua McMichael, Adam C. Coffman, Susanna Kiwala, Kilannin Krysiak, Alex H. Wagner, Malachi Griffith, Obi L. Griffith

## Abstract

**Summary:** The druggable genome encompasses the genes that are known or predicted to interact with drugs. The Drug-Gene Interaction Database (DGIdb) provides an integrated resource for discovering and contextualizing these interactions, supporting a broad range of research and clinical applications. DGIdb is currently accessed through structured web interfaces and API calls, requiring users to translate natural-language questions into database-specific query patterns. To allow for the use of DGIdb through natural language, we developed the DGIdb Model Context Protocol (MCP) server, which allows large language models (LLMs) access to up-to-date information through the DGIdb API. We demonstrate that the MCP server greatly enhances an LLM’s ability to answer questions requiring accurate, up-to-date biomedical knowledge drawn from structured external resources.

**Availability and implementation:** The DGIdb MCP server is detailed at https://github.com/griffithlab/dgidb-mcp-server and includes instructions for accessing the server through the Claude desktop app.

## 1. Introduction

Drug-gene interactions play a central role in precision medicine, as they help determine which treatments are likely to be effective for a given patient and which may lead to resistance or adverse effects. As large-scale molecular studies increasingly reveal large sets of genes associated with disease, researchers need tools that can connect these findings to potential therapeutic strategies. The Drug-Gene Interaction Database (DGIdb)^1–5^ addresses this need by aggregating and harmonizing drug-gene interaction information from multiple sources, providing a centralized resource for assessing gene druggability and therapeutic relevance. DGIdb supports downstream research and clinical-informatics applications, including gene-drug linking for prioritized disease genes^6^, exploratory hypothesis generation^7^, drug repurposing^8^, genomic gene-list annotation^9^, and precision oncology workflows^10^. The value of searching a catalog of drug-gene interactions is well illustrated by the challenge of *FLT3-mutated Acute Myeloid Leukemia*. Although the FLT3 inhibitor *Gilteritinib* is initially effective, patients often develop resistance due to additional genetic changes that emerge during treatment.^11^ In such settings, DGIdb could be used to identify drugs known or predicted to interact with genes implicated in resistance, supporting exploratory analyses and prioritization of alternative therapeutic candidates.

Despite DGIdb’s utility, this process can be time-consuming in multistep analyses, such as studies of acquired drug resistance, where users must first identify resistance-associated genes, then perform separate queries for each gene, and finally manually combine and prioritize potentially large lists of interacting drugs. As a result, DGIdb is difficult to use directly within natural-language workflows and is largely inaccessible to large language models (LLMs), which cannot natively construct or execute these structured queries. At the same time, LLMs have emerged as promising tools for a wide range of biomedical applications, including literature summarization and clinical decision support, due to their ability to synthesize complex information and respond flexibly to natural-language questions.^12–16^ However, LLMs rely on static internal knowledge, may generate inaccurate or unsupported statements, and cannot directly access curated biomedical resources without additional infrastructure.^17^ These limitations restrict their reliability in settings where accuracy and traceability are essential.

To address these challenges, the Model Context Protocol (MCP) provides a standardized way to connect LLMs to authoritative external data sources. MCP servers serve as a bridge between the flexibility of LLMs and the precision of curated biomedical databases and have been increasingly applied to biomedical domain APIs, databases, and literature search endpoints, including variant-level knowledgebases via the CIViC MCP server.^18–20^ The DGIdb MCP server builds upon this framework, allowing LLMs and researchers to retrieve, interpret, and cite drug-gene interaction information directly from the curated resources, making it easier to integrate accurate interaction data into genome-wide analyses and therapeutic decision-making.

### 2. DGIdb MCP Server Description

The DGIdb MCP server is publicly available through Cloudflare. The server provides four tools, each corresponding to predefined GraphQL queries for integration with LLMs. These tools allow LLMs to retrieve drug information, gene information, and drug-gene interactions based on either a supplied gene list or drug list. The drug information tool returns key attributes such as FDA approval status, drug classifications including immunotherapy and antineoplastic drugs, and additional characteristics such as drug class. The gene information tool provides DGIdb gene category annotations, such as kinase, ligase, and nuclear receptor designations, along with other attributes relevant to gene druggability. Together, these data sources supply structured and well-curated information that supplements an LLM’s understanding of the drug-gene interaction space. When the LLM calls the corresponding interaction query with a supplied list of genes or drugs, the server returns a ranked list of their known interactions. Each interaction includes an interaction type and, when applicable, directionality (activating or inhibitory) which together describe the mechanism of action and effect on biological activity. Every interaction is also linked to its supporting evidence, which may come from a publication or a curated database. Interactions are ranked primarily according to the FDA approval status of the associated drugs, and secondarily according to the DGIdb interaction score, which was introduced in the DGIdb 4.0 release.^4^ The score combines evidence strength (number of supporting publications and curated sources) with interaction specificity, calculated from the relative number of known partners for the drug and gene compared to global averages. The final score prioritizes interactions that are both well supported and more specific, and remains static within a data release. This ranking strategy allows LLMs to prioritize clinically informative and well-supported interactions.

To accommodate spelling variations and alternate names, the MCP server normalizes drug and gene inputs to the closest matching label recognized by DGIdb. Alias similarity is scored using Dice’s coefficient over bigrams, which requires limited memory making it ideal for Cloudflare workers. The MCP server obtains drug and gene aliases from the VICC Normalization Services, which provide a standardized set of identifiers and synonyms for use during the scoring process.^21,22^

### 3. Example Use Cases

1. A user might ask, “Which FDA-approved drugs target the gene *KIT*, and what is the supporting evidence?” In response, the LLM identifies the gene of interest and the user’s intent, then invokes the appropriate MCP tool. The MCP server normalizes the gene name and issues the corresponding GraphQL query to the DGIdb API. It returns a ranked list of drugs that interact with *KIT*, along with interaction metadata such as interaction score, interaction type and direction, FDA-approval status, and supporting sources. Using this structured output, the LLM composes a user-facing answer that includes relevant evidence and links to the cited publications. A more complete list of use cases for the DGIdb MCP server are outlined in Supplementary Table 1.
2. A second example use case demonstrates the joint use of the CIViC MCP^20^ and DGIdb MCP servers for drug candidate selection (Figure 1). A user may ask, “What genes can cause resistance to *Ibrutinib* in *Chronic Lymphocytic Leukemia*, and what alternative drugs can target them?” The LLM identifies the relevant CIViC entities (Therapy, Disease, Clinical Significance) and invokes the appropriate tool from the CIViC MCP server to retrieve variant evidence. The CIViC MCP server issues a GraphQL query to the CIViC API, which returns structured Evidence Items associated with *Ibrutinib* resistance in *CLL*. From these records, the LLM extracts *BTK* as the primary gene implicated in resistance and composes a query to the DGIdb MCP server to identify other drugs that interact with *BTK*. The server returns a ranked list of drug-gene interactions, enabling the LLM to respond with the highest-priority drug candidates, including *Tirabrutinib, Acalabrutinib*, and *Zanubrutinib*. Using the outputs from each MCP server, the LLM returns an answer that cites relevant curated evidence and publications. Additional examples of how the DGIdb and CIViC MCP servers can be used together are outlined in Supplementary Table 2.

**Figure 1.**
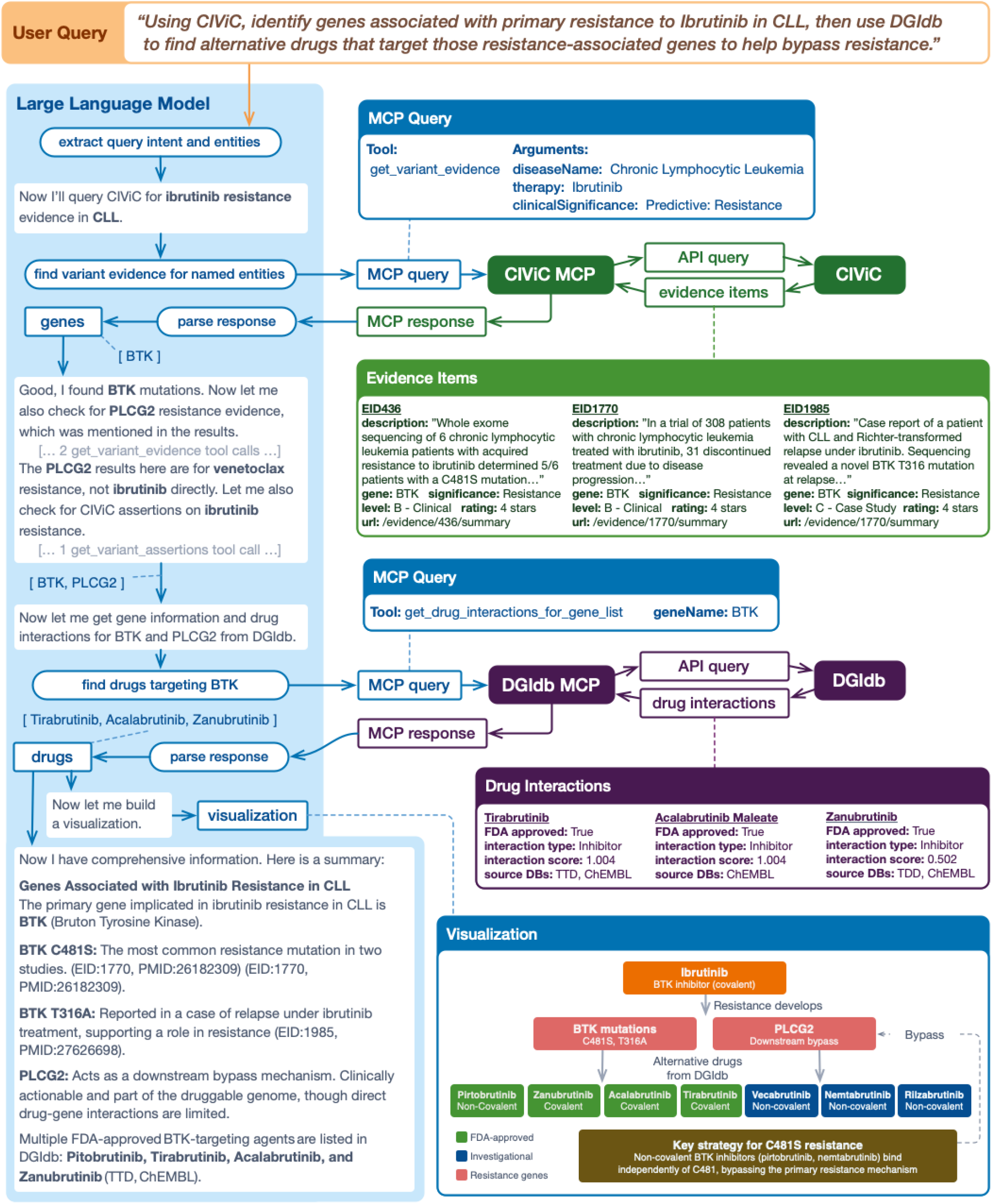
Joint use of the CIViC MCP and DGIdb MCP servers for drug candidate selection with Claude. A user asks, “What genes can cause resistance to Ibrutinib in Chronic Lymphocytic Leukemia, and what alternative drugs can target them?” The LLM: (1) extracts the users intent and relevant entities (Therapy, Disease, Significance), (2) composes a query to the CIViC MCP server with the appropriate tool selection, (3) the CIViC MCP server queries the CIViC API and returns Evidence Items, (4) the LLM parses the MCP response and identifies the BTK as an ibrutinib resistance gene, (5) the LLM composes a new query to the DGIdb MCP server to identify other drugs that interact with BTK, (6) the DGIdb MCP server queries the DGIdb API and returns drug interactions for BTK, (7) the LLM composes a summary.

### 4. Benchmarking LLM Performance with DGIdb and CIViC MCP Servers

We developed a set of classification and information-retrieval tasks to measure how effectively the MCP server enhances an LLM’s ability to answer questions requiring accurate and current biomedical knowledge. As a first task, we asked GPT-5 to label drugs according to three DGIdb-derived attributes: FDA approval status, immunotherapy classification, and antineoplastic classification (Supplementary Table 3). We selected 100 drug names using stratified sampling to approximate a 50/50 balance of positive and negative examples for each label. Because relatively few drugs in DGIdb are classified as immunotherapies, achieving a perfectly even split was not possible. GPT-5 was prompted to return three comma-separated YES/NO values to facilitate parsing (Supplementary Prompt 1). Across all labels, GPT-5 augmented with the DGIdb MCP substantially outperformed GPT-5 alone achieving a macro weighted F1-score of 0.99 compared to 0.75. GPT-5 without MCP struggled particularly with identifying immunotherapies, producing many false negatives and yielding a recall of only 0.38.

Next, we compared how often GPT-5 would invoke the DGIdb MCP server when requesting the same drug information without explicitly mentioning DGIdb (Supplementary Prompt 2). The DGIdb MCP server was available to the model in the same configuration as the previous task, but tool use was not explicitly instructed. GPT-5 did not use the DGIdb MCP server for 55 out of the 100 drugs. Among cases where GPT-5 did not invoke the MCP server, performance declined, particularly for immunotherapy and antineoplastic classification. Recall dropped from 1.00 to 0.68 for immunotherapies and from 0.88 to 0.68 for antineoplastic drugs (Supplementary Table 4). To understand when GPT-5 decided to use the MCP server, we manually inspected a random subset of predictions. In these examples, the MCP server was generally not invoked for widely used, FDA-approved therapeutics with unambiguous, commonly referenced names (e.g. Tamoxifen), but was preferentially used for uncommon, structurally complex molecules, or database-style identifiers (e.g. ODM-203), indicating that GPT-5 selectively employed the MCP server when additional disambiguation or external grounding was likely required.

Lastly, we evaluated a task that used both the CIViC MCP and DGIdb MCP servers to demonstrate the potential of LLM-assisted drug candidate selection, as outlined in Supplementary Figure 1. To allow for direct evaluation of the intermediate gene list, this task utilized two prompts. In the first step, GPT-5 was prompted to identify genes associated with resistance to a specified therapy in a given disease using CIViC-derived information (Supplementary Prompt 3). In the second step, we issued a separate prompt for each gene identified in the first step, asking GPT-5 to list drugs that interact with that gene using DGIdb-derived information (Supplementary Prompt 4). To keep runtime manageable, we limited the request to the top 20 interacting drugs per gene, as full interaction lists for a single gene can exceed 500 entries. For this evaluation, we randomly selected 50 disease-therapy combinations from CIViC evidence as starting points for the drug-candidate identification process. GPT-5 was instructed to rank genes according to the strength of their associated CIViC evidence. Genes were ordered first by Evidence Level, with level A considered the strongest and level E the weakest, and then by Evidence Rating, prioritizing higher ratings (5 to 1). Drugs were ranked primarily by FDA approval status and secondarily by interaction score. Because the multihop task requires GPT-5 to return genes and drugs in a prioritized order, we evaluate correctness using precision, recall, and F1, and we assess ranking quality using normalized discounted cumulative gain (NDCG). NDCG is a common information retrieval metric that measures how well the model ranks relevant items, giving greater weight to items that appear earlier in the list. NDCG ranges from 0 to 1, with higher values indicating rankings that more closely match the ideal ordering. GPT-5 augmented with the CIViC and DGIdb MCP servers substantially outperformed GPT-5 alone on the drug-list task, improving F1 from 0.14 to 0.95 and NDCG from 0.19 to 0.93 (Supplementary Table 5).

## 5. Conclusion

We demonstrate that the DGIdb MCP server enables reliable, structured exploration of drug-gene interactions by allowing an LLM to query a comprehensive, curated interaction resource in response to natural-language questions. By grounding GPT-5 in DGIdb, this approach supports accurate retrieval, ranking, and contextualization of interaction evidence, which is particularly important for tasks such as identifying alternative therapeutic candidates and interpreting resistance-associated genes.

We further show that LLMs can make effective use of multiple MCP servers simultaneously. In a drug-candidate identification task for therapy resistance, GPT-5 successfully combined variant-level evidence retrieved from the CIViC MCP server with drug-gene interaction data from the DGIdb MCP server, demonstrating how MCP servers can be composed to support multi-step biomedical reasoning workflows that require information from multiple sources.

Our evaluation also highlights an important practical consideration: GPT-5 did not consistently invoke MCP servers unless the relevant biomedical resource is mentioned explicitly in the prompt. When the source (e.g., “DGIdb”) is named, GPT-5 reliably delegates retrieval to the MCP server. When it is omitted, however, the model often defaults to its internal knowledge, resulting in reduced accuracy, particularly for identifying immunotherapies. This finding emphasizes the importance of prompt design in ensuring consistent MCP usage.

While the MCP framework has previously been applied to variant-level knowledge through the CIViC MCP server^20^, this work extends the paradigm to the drug-gene interaction domain and illustrates how multiple MCP servers can be integrated to address more complex biomedical questions. As future work, the development of standardized benchmarks for MCP-assisted biomedical tasks, such as drug-gene interaction retrieval, variant interpretation, and cross-knowledgebase reasoning, will be critical for enabling consistent evaluation and comparison of MCP-enabled LLM systems.

