## Supplemental Materials for "Searching the Druggable Genome using Large Language Models"

**Table 1: DGIdb MCP Example Use Cases**

| **Category** | **Example Query** | **Intended Output*** |
| --- | --- | --- |
| Drug Discovery | Which FDA-approved drugs target the gene KIT? | A ranked list of FDA-approved KIT-interacting drugs with interaction scores, interaction types, approval status, and links to supporting sources. [Chat Link](https://claude.ai/share/62d8e788-7b0f-453a-af08-dc066d34b9b1). |
| Drug Info | What information is available about Imatinib? | A structured overview of Imatinib including its FDA approval status, year of approval, therapeutic and drug class annotations. [Chat Link](https://claude.ai/share/999dfa2e-aa4b-4a5c-bcf7-9be72886b873). |
| Gene Category Info | What categories and functional annotations are associated with the gene BRAF? | A structured summary of BRAF annotations, including clinical actionability, drug resistance relevance, druggability, enzyme class, and kinase classification, with the supporting source databases listed for each category. [Chat Link](https://claude.ai/share/91a62f7d-d278-4a4b-a70c-8f54d598d147). |
| Gene Symbol Disambiguation | Is FLT3 an unambiguous gene symbol? | An assessment of whether FLT3 maps uniquely to a single gene, including any known aliases/synonyms, overlapping symbols, and the resolved canonical gene identifier(s), with supporting source links. [Chat Link](https://claude.ai/share/5fa0883b-538b-47c8-8b68-99b8d9db4045). |
| Mechanism-specific targeting | Which inhibitors interact with EGFR? | A filtered list of EGFR inhibitors, prioritized by FDA approval and interaction score, with evidence sources cited. [Chat Link](https://claude.ai/share/0daf9c70-b6cd-4c00-b2af-ff2c91a6aff7). |
| Interaction type interpretation | What genes interact with the drug Ibrutinib and by what mechanism of interaction? | A structured summary of Ibrutinib-gene interactions, annotated with interaction direction and type (e.g., inhibitor, modulator). [Chat Link](https://claude.ai/share/1297b4d8-d55b-408d-9648-ef34e266f7ca). |

Chat links to conversations with Opus 4.5 via Claude Desktop on 1/14/2026.

*API responses are not visible in the linked chats.

**Table 2: DGIdb MCP + CIViC MCP Use Cases.**

| **Category** | **Example Query** | **Intended Output*** |
| --- | --- | --- |
| Resistance-guided drug discovery  (CIViC→DGIdb) | What genes can cause resistance to ibrutinib in chronic lymphocytic leukemia, and what alternative drugs can target them? | CIViC-supported resistance genes for ibrutinib in CLL and a ranked list of interacting alternative drugs from DGIdb, with citations. [Chat Link](https://claude.ai/share/be18a9b9-0a1b-4831-9dc9-9f72dd9d5049). |
| Oncogenicity-driven functional validation  (CIViC→DGIdb) | What is the evidence for oncogenicity of variants in ERBB2 for breast cancer, and what inhibitors could I use to experimentally validate their importance in vitro? | CIViC-supported oncogenic ERBB2 variants in breast cancer with summarized evidence and citations, plus a ranked list of ERBB2-targeting inhibitors from DGIdb suitable for in vitro validation. [Chat Link](https://claude.ai/share/1bd81002-fe12-44aa-9b5c-775a0fb426d2). |
| Drug target profiling with clinical evidence  (DGIdb→CIViC) | What genes interact with the drug Ibrutinib, and which of those genes have specific variants in CIViC associated with sensitivity or resistance? | A list of genes interacting with ibrutinib annotated with CIViC sensitivity or resistance evidence where available. [Chat Link](https://claude.ai/share/9ad52a71-74c1-40a5-aa22-afc03e5fa940). |
| Target resistance interpretation (DGIdb→CIViC) | Are there known resistance mechanisms for genes targeted by Osimertinib? | Osimertinib target genes with CIViC-documented resistance mechanisms and supporting evidence links. [Chat Link](https://claude.ai/share/9da56313-74b4-4978-954b-43437a63e47f). |
| Gene-guided therapy matching  (DGIdb→CIViC) | Which drugs interact with PIK3CA, and is there evidence of sensitivity to these drugs in breast cancer? | PIK3CA-interacting drugs prioritized by DGIdb and annotated with CIViC sensitivity evidence in breast cancer. [Chat Link](https://claude.ai/share/b73748ee-74d6-4b7f-92f1-8a8620960bcd). |
| Expression-based therapy matching  (DGIdb→CIViC) | For tumors with EGFR amplification, which drugs target EGFR and is there clinical evidence supporting sensitivity? | EGFR-targeting drugs with CIViC evidence supporting sensitivity in EGFR-amplified tumors. [Chat Link](https://claude.ai/share/b2d6ffa7-a2a2-4c2f-9805-3d6aad577362). |

Chat links to conversations with Opus 4.5 via Claude Desktop on 1/14/2026 and 3/25/2026.

*API responses are not visible in the linked chats.

**Table 3.** **Drug classification performance.** N represents the number of positive samples for each label.

|  | **GPT-5 + DGIdb MCP** | | | **GPT-5** | | |
| --- | --- | --- | --- | --- | --- | --- |
| **Label** | **Precision** | **Recall** | **F1-score** | **Precision** | **Recall** | **F1-score** |
| FDA-Approved (N=48) | 1.00 | 0.98 | 0.99 | 0.89 | 1.00 | 0.94 |
| Immunotherapy (N=47) | 1.00 | 1.00 | 1.00 | 0.95 | 0.38 | 0.55 |
| Antineoplastic (N=64) | 1.00 | 1.00 | 1.00 | 0.86 | 0.69 | 0.77 |

**Table 4. Drug classification performance.** GPT-5 with access to the DGIdb MCP server but with no mention of DGIdb in the prompt. N represents the number of positive samples for each label.

|  | **MCP Used (N=45)** | | | **No MCP Usage (N=55)** | | |
| --- | --- | --- | --- | --- | --- | --- |
| **Label** | **Precision** | **Recall** | **F1-score** | **Precision** | **Recall** | **F1-score** |
| FDA-Approved (N=48) | 0.91 | 1.00 | 0.95 | 0.93 | 1.00 | 0.96 |
| Immunotherapy  (N=47) | 1.00 | 1.00 | 1.00 | 0.95 | 0.68 | 0.80 |
| Antineoplastic (N=64) | 1.00 | 0.88 | 0.94 | 0.91 | 0.68 | 0.78 |

**Randomly sampled examples:**

**No MCP Usage:** TAMOXIFEN, TREMELIMUMAB, RITUXIMAB, NIVOLUMAB, PEGINTERFERON ALFA-2A, [125I]TYR45-KISSPEPTIN-15, [D-TYR8]CYN 154806, DULANERMIN, ENTRECTINIB, GOLIMUMAB

**MCP used:** ODM-203, LBY-135, ISOBAVACHALCONE, INTERFERON ALFA-N3, CHEMBL_CHEMBL19954, CIXUTUMUMAB, ABT-239, (+)-ADTN, CS+, JNK INHIBITOR 9L

**Table 5. Drug Candidate Identification Performance.**

| **Model** | **Metric** | **CIViC Gene List** | **DGIdb Drug List** |
| --- | --- | --- | --- |
| **GPT-5 + MCP Servers** | Precision | 0.96 | 0.96 |
|  | Recall | 0.96 | 0.95 |
|  | F1-score | 0.96 | 0.95 |
|  | NDCG | 0.97 | 0.93 |
| **GPT-5** | Precision | 0.23 | 0.23 |
|  | Recall | 0.43 | 0.16 |
|  | F1-score | 0.30 | 0.14 |
|  | NDCG | 0.44 | 0.19 |

**Prompt 1: DGIdb Drug Info**

**System:** "You are a chatbot for the Drug Gene Interaction Database (DGIdb)"

**User:** “Drug: {drug_name}

For the drug named above ONLY, state if each property applies (YES/NO) based on DGIdb.

Output EXACTLY this 2-line CSV (no extra text):

fda_approval,immunotherapy,anti_neoplastic

<YES/NO>,<YES/NO>,<YES/NO>”

**Prompt 2: DGIdb Drug Info w/ MCP, no mention of DGIdb**

**System:** “You are a helpful AI assistant with access to external tools. Follow the user's instructions carefully.”

**User:** “Drug: {drug_name}

For the drug named above ONLY, state if each property applies (YES/NO).

Output EXACTLY this 2-line CSV (no extra text):

fda_approval,immunotherapy,anti_neoplastic

<YES/NO>,<YES/NO>,<YES/NO>”

**Prompt 3: CIViC Genes Responsible For Resistance**

**System:** "You are a chatbot for CIViC and the Drug Gene Interaction Database (DGIdb). "

When listing genes from CIViC, you must rank them using CIViC evidence strength."

GENE RANKING RULES:

1. Rank genes first by evidenceLevel (A strongest → E weakest).

2. Within the same evidenceLevel, rank genes by evidenceRating (5 → 1 → null).

3. Return only HGNC gene symbols, in this ranked order, as a comma-separated list

like: GENE1,GENE2,GENE3."

**User: “**According to CIViC, list genes whose variants/molecular profiles are associated with resistance to {therapies} in {disease_name} (significance = RESISTANCE)”

**Prompt 4: DGIdb Candidate Drugs**

**System:** “You are a chatbot for the CIViC and Drug Gene Interaction Database (DGIdb). You receive a gene and DGIdb interaction for that gene. Your task is to return a ranked list of up to 20 drugs.

RANKING RULES:

1. List FDA-approved drugs first.

2. Within the approved drugs, sort by the DGIdb interactionScore in descending order

3. After all approved drugs, list non-approved drugs, also sorted by interactionScore in descending order.

4. Return only drug names (no scores), as a comma-separated list like: DRUG1,DRUG2.”

**User:** “What drugs target or modulate this gene product in DGIdb? Gene: {gene}

Return a comma-separated list of up to 20 drug names following the ranking rules.”

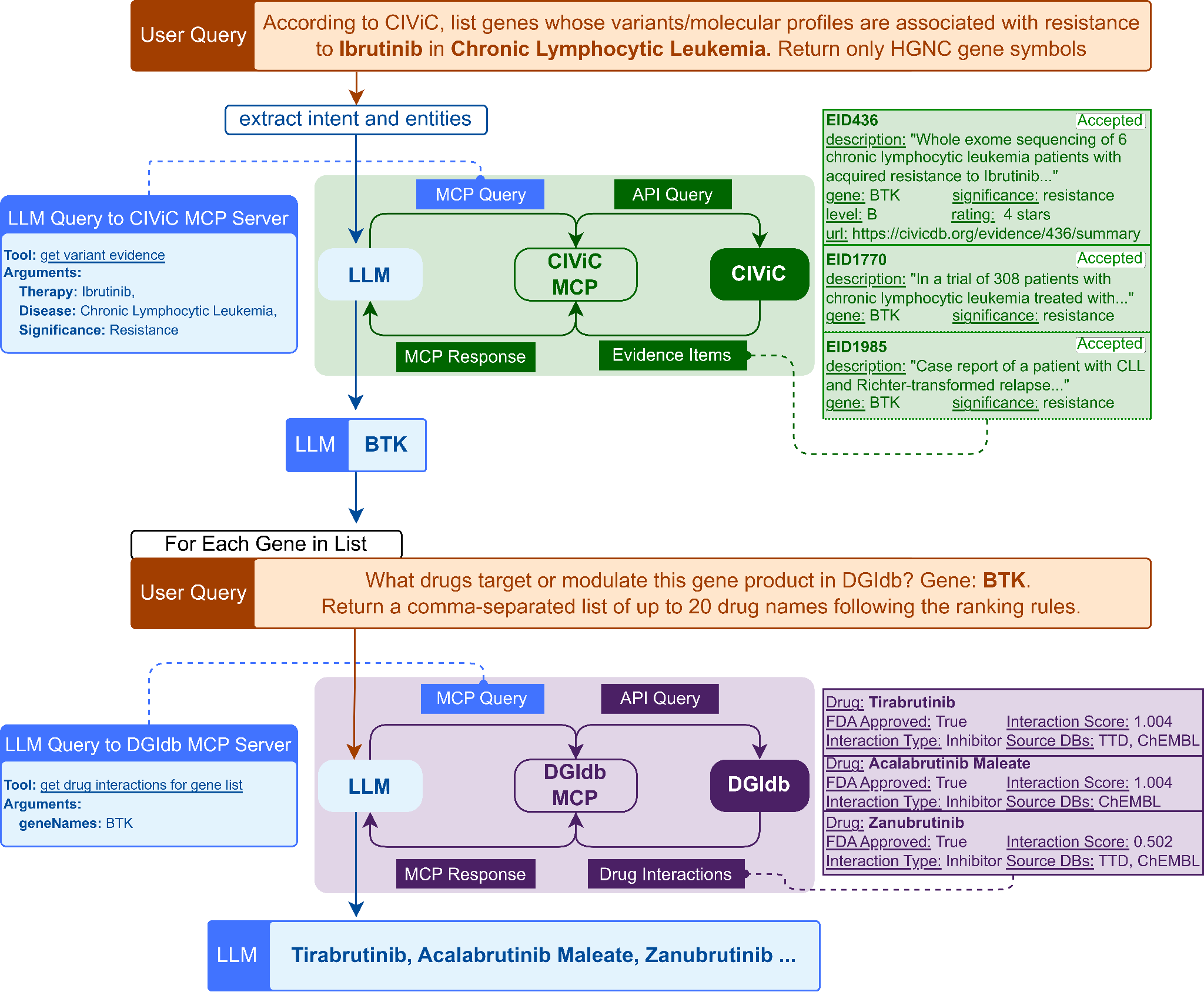

**Figure 1.** **Evaluation workflow overview.** The LLM is first prompted to provide the list of ordered genes. Then individually prompted to provide the list of interacting drugs for each gene.
